# Tools for Quantitative Analysis of Calcium Signaling Data Using Jupyter-Lab Notebooks

**DOI:** 10.1101/2023.06.13.544740

**Authors:** John Rugis, James Chaffer, James Sneyd, David Yule

## Abstract

Calcium signaling data analysis has become increasing complex as the size of acquired datasets increases. In this paper we present a Ca^2+^ signaling data analysis method that employs custom written software scripts deployed in a collection of Jupyter-Lab “notebooks” which were designed to cope with this complexity. The notebook contents are organized to optimize data analysis workflow and efficiency. The method is demonstrated through application to several different Ca^2+^ signaling experiment types.

## 1. Introduction

In a biological research laboratory context, data analysis requires transforming experimental measurements into a mechanistic understanding of an event. This transformation process is invariably computer based and has become increasingly complex, working with ever larger quantities of data. Not only does the analysis process have to accommodate large amounts of data, but there is also the challenge of moving from a qualitative to a quantitative interpretation.

This is especially true of large data sets acquired from Ca^2+^ signaling experiments. Often contemporary imaging methods are designed to acquire information with high temporal and spatial fidelity from numerous individual cells and/or from multiple intracellular regions of interest. Interrogating these increasingly large data sets to provide quantitative information relating to latency, magnitude of response, frequency of events and spatial homogeneity is required to yield mechanistic insight into cellular processes controlling Ca^2+^ homeostasis and is increasingly labor intensive. In our experience, processing methods for Ca^2+^ signaling datasets share some common requirements but can also be highly specific to the needs of a particular lab and no current off-the-shelf software tool meets all these needs. (See the discussion section for further details about this.) The overarching goal is quantifiable, consistent, repeatable analysis not only within a given research group, but between different groups within the research community.

To fulfill these requirements, we generated custom Python [1, 2] computer scripts which are deployed in a collection of Jupyter “notebooks” [3]. These software tools facilitate calculation of a number of common parameters from Ca^2+^ signaling data sets. The utility and flexibility of this framework has addressed the needs of our research program and we envision that it will have utility for other users as well. These notebooks were designed to gather and compare data relating to the amplitude and frequency of responses from either *in vivo* or *in vitro* experiments.

Our notebook collection is organized into two general categories: main processing and optional post-processing. Main processing consists of region-of-interest (ROI) identification and post-processing includes notebooks for signal peak identification, frequency analysis and movie making. Detected ROIs are used to isolate and extract subsets of image data that are associated with individual cells under study. Peak analysis, in various forms, is used to quantify Ca^2+^ signal responses.

Our notebooks have been designed to automate repetitive steps and offer users the flexibility of diagnostic interactive data exploration. Notebook script features include generating plots, saving raw summary data in comma-separated-values (csv) files, and creating time-stamped directories for organizing results.

## 2. Methods

We have developed a collection of eight notebooks, namely:

1. ROI_Detection
2. Ratiometric_ROI_Detection
3. Peak_Detection
4. CSV_Peak_Detection
5. CSV_Peak_Detection_SOCE
6. CSV_Peak_Detection_Inhibitor
7. Frequency_Analysis
8. Movie_Making

These notebooks provide a flexible collection of analysis tools which can be applied to a broad range of Ca^2+^ signaling experiments. Each experiment in our laboratory uses two or more of these notebooks for data analysis.

The notebooks employ a number of numerical methods, many of which originate from within computational computer graphics. For example, Canny edge detection [4] is used to locate the outlines of individual cells in the microscopy images under study. Flood fill [5] is used to graphically “fill in” outlined regions in images. The standard graphics morphology operations image erosion and dilation [6] are used to “shrink” and “grow” the size of identified image regions. Watershed image segmentation [7] is used to separate overlapping regions in images.

## 3. Experiments and Results

The experiments in our laboratory fall broadly onto four types: single wavelength indicator experiments, experiments that quantify agonist induced Ca^2+^ signaling, experiments that quantify store-operated calcium entry (SOCE), and experiments to define the efficacy of inhibitors of Ca^2+^ signaling pathways. Note that, with time varying fluorescence microscopy imaging of biological samples, it is essential to be able isolate specific cellular regions-of-interest in the image stack sequence for further analysis and this is a common requirement in each of our experiment types. Automated region of interest (ROI) detection is a key feature of our notebooks. This process was formerly highly labor-intensive and time consuming.

This section gives an overview of how we employed the notebooks to analyze data from each of our different experiment types. A more detailed step-by-step outline of notebook usage is given in the supplemental notes that can be found at the link: https://github.com/jrugis/cell_tools/blob/master/Supplemental_Notes.pdf

### 3.1 Single Wavelength Indicator Experiments

A wide variety of so-called “single wavelength” or “single excitation” Ca^2+^ indicators are now commonly used [8]. These probes exhibit fluorescence properties providing excitation and emission characteristics throughout the visible spectrum. Both chemical and genetically encoded probes [9] are available which exhibit high quantum efficiency, have high dynamic range and Ca^2+^ affinities useful for probing both bulk cytosolic and local [Ca^2+^]. Next, we describe a series of notebooks designed to analyze and present data generated using a single excitation wavelength indicator.

#### 3.1.1. Example: gCamp Experiment

The analysis of gCamp experiments requires the use of the ROI_Detection, Peak_Detection and optionally the Frequency_Analysis and Movie_Making notebooks. Although the notebooks are described using the Mistgcamp-3 dataset, user-determined parameters offer flexibility in the types of single wavelength indicator data that can be processed.

The data in this experiment were generated from mice expressing the genetically encoded Ca^2+^ indicator gCamp6F, specifically in exocrine cells [10, 11]. The signals were recorded by multiphoton microscopy in live mice and the spatial heterogeneity of the Ca^2+^ signal necessitated a means of automating ROI selection. In this example, microscopy image data is stored in files each of which contain a time series image stack associated with a particular experimental sample stimulus frequency. Analysis generally consists of comparing the effects in a field of view of increasing stimulation frequencies, analogous to a stimulus vs response experiment performed in vitro. This paradigm produced multiple files to be subjected to analysis.

Region-of-interest detection begins with generating an initial unstimulated reference “ground state” image which is a per pixel average (or standard deviation) over the time period in which the sample is not stimulated. This is followed by generating a “stimulated time” image, which is again a per pixel average (or standard deviation) over the time during which the sample is stimulated. Then a difference image is created by doing a per-pixel subtraction of the unstimulated image from the stimulated image. Note that this has the effect of removing background structure and spurious bright areas, primarily leaving active cell apical regions as can be seen in **Figure 1**.

**Figure 1.**
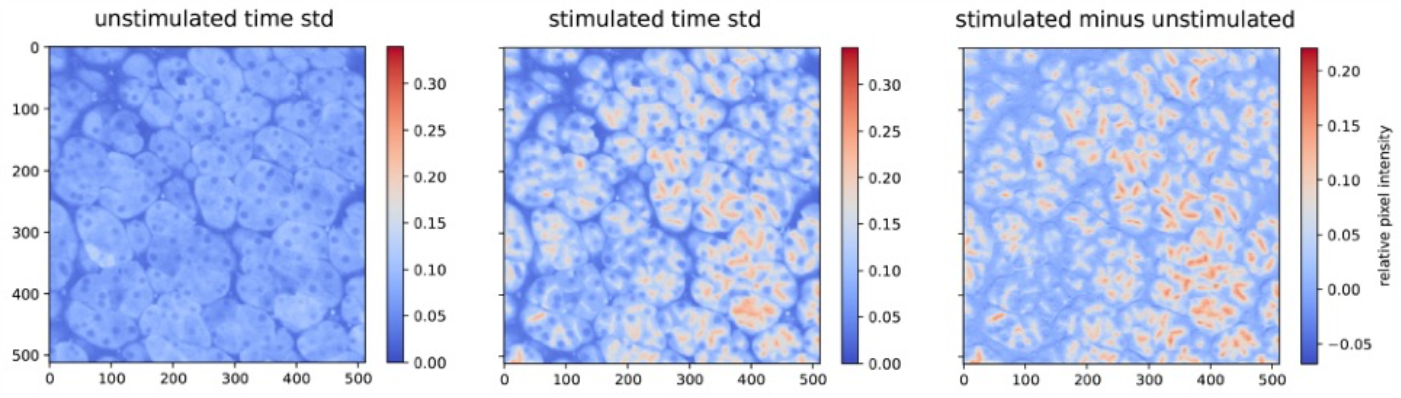
Stimulated, unstimulated and difference images of pixel intensities from the 5Hz Mistgcamp-3 dataset.

Next, cell apical regions are individually identified using simple pixel intensity thresholding. Subsequent filtering removes undesired small regions and smooths the remaining regions giving the final. Further details of the ROI identification process for this experiment can be found in the supplemental notes.

Analysis continues using the Peak_Detection notebook. The design goal with this notebook was to characterize biological response plot curves in a way that is both intuitively meaningful to physiologists and clearly quantifiable. The characterization was designed to match practitioner consensus on identifying the most prominent peaks using a mathematically well-defined algorithm which, in that sense, removes subjective bias. Due to the anticipated wide variation in signal plots, the peak detection notebook does include a limited number of user-adjustable parameter settings.

Detected peaks for each stimulation frequency are plotted in the notebook as black dots overlaid on region summary plots as shown, from the Mistgcamp-3 dataset, in **Figure 2Figure 2**. Additionally, each plot trace is analyzed to determine the time of stimulation onset (indicated using red dots in the figure), peak counts, area under the curve, first peak height, and maximum peak height. Latency, risetime and maximum slope from stimulation onset to first peak are also calculated.

**Figure 2.**
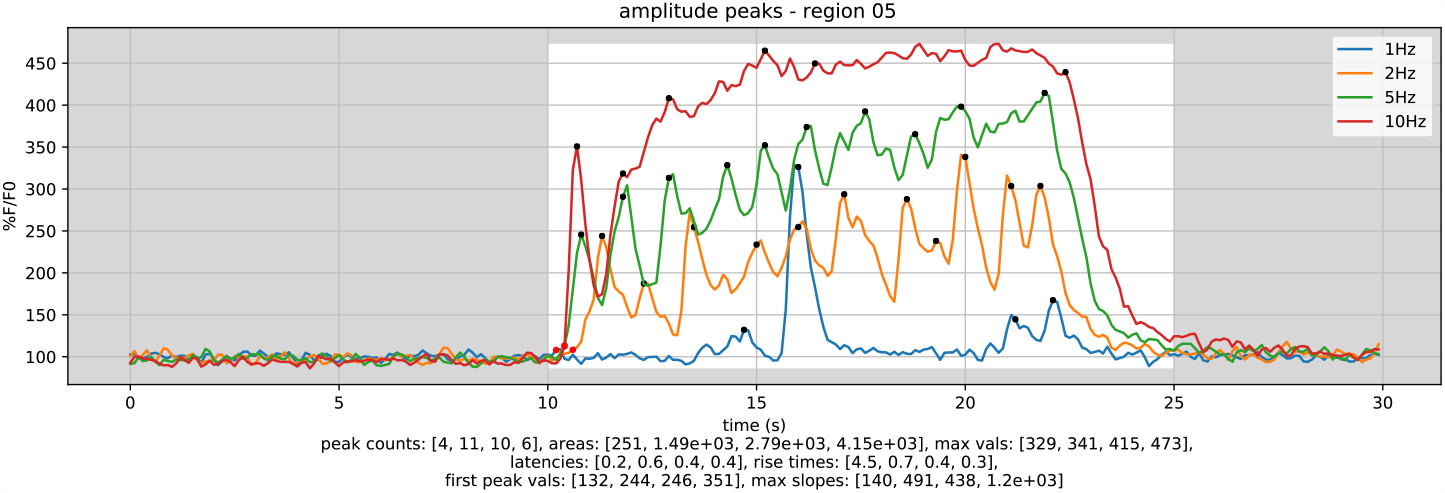
Peaks summary plot for four different stimulation frequencies within one region in the Mistgcamp-3 dataset. Plot trace numerical analysis results are shown below the plot.

### 3.2 Agonist-induced Ca^2+^ Signaling Experiments

A host of hormones, neurotransmitters and growth factors signaling through phospholipase C lead to an increase in cytosolic [Ca^2+^] [12, 13]. A common and laborious experimental task following imaging experiments is to quantify these changes in [Ca^2+^], often from numerous cells in a field of view and captured over time with fast frame rates. Here we describe notebooks to first assign regions of interest (ROIs) to cells and subsequently to quantify various parameters associated with increases in [Ca^2+^].

#### 3.2.1 Example: CCh and Trypsin Experiments

Carbachol (CCh) and trypsin bind to muscarinic and protease-activated receptors, respectively, couple through Gαq to cause an increase in inositol 1,4,5-trisphosphate concentration and thus induce Ca^2+^ release from the endoplasmic reticulum (ER) into the cytosol of the cell [12]. The [Ca^2+^] is commonly measured by changes in the ratiometric indicator, Fura-2 [14, 15]. The concentration vs. response relationship and the characteristics of the [Ca^2+^] signal differ between stimulation with CCh or trypsin. The analysis of both the CCh and trypsin experiments is facilitated using the Ratiometric_ROI_Detection and CSV_Peak_Detection notebooks.

The Ratiometric_ROI_Detection notebook has been customized to work specifically with imaging data collected using ratiometric indicators exemplified by Fura-2. This notebook features automation which allows multiple experiments to be analyzed simultaneously. The notebook takes as input both the image stacks generated by excitation at 340 nm and the 340/380 nm ratio time series image stacks for each experiment being analyzed.

The CSV_Peak_Detection notebook is used to quantitatively analyze the csv files output by the Ratiometric_ROI_Detection notebook (or manually created csv files in the same data format). The CSV_Peak_Detection notebook features batch analysis of typical agonist-induced Ca^2+^ signal data from each of the cells identified by the ROI identification process.

Configuration settings within the CSV_Peak_Detection notebook include specifying one or more time zones indicating the time period in which the cells under study were stimulated. The results of peak detection and analysis include latency, slope, and rise time associated with the first peak (red color) in each stimulation time zone (white background color), for example, in **Figure 3**.

**Figure 3.**
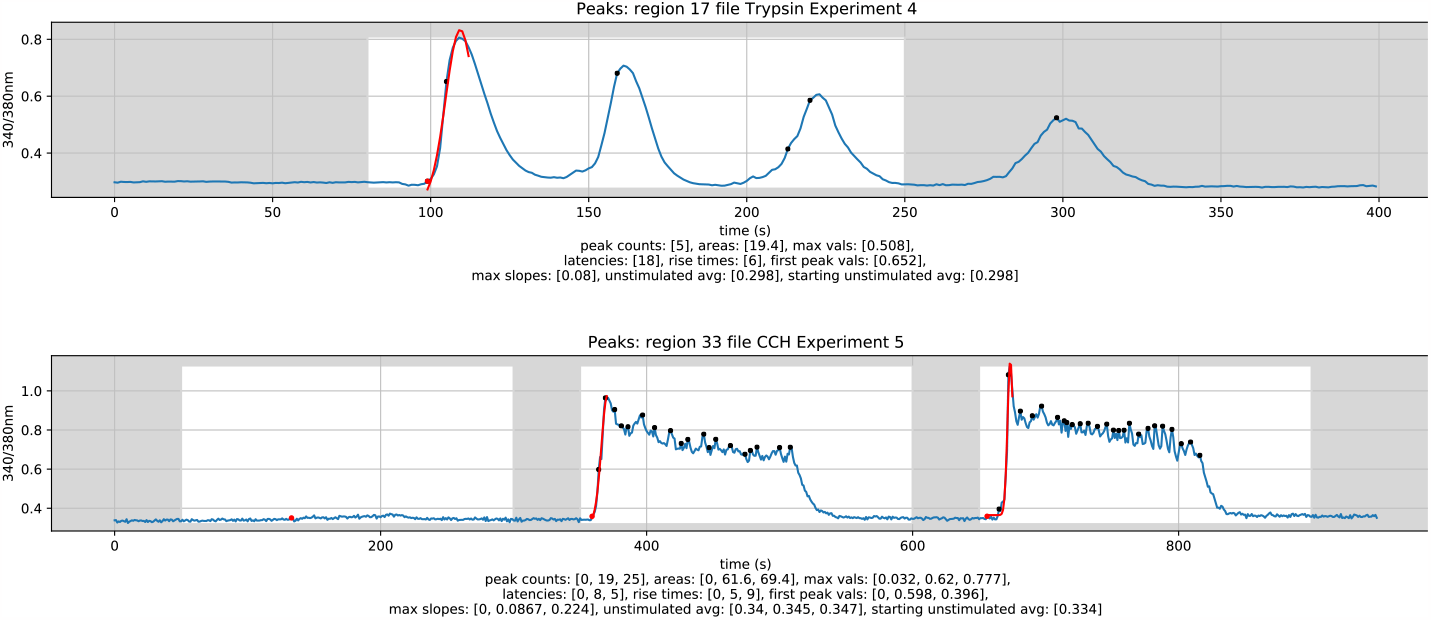
Example analysed plot output from Trypsin Experiment 4 Region 17 (top) and CCh Experiment 5 Region 33 (bottom)

Analysis results (csv files) and plots (pdf files) are automatically saved. Each of the values contained within a “total_data_by_zone” csv file are useful for quantifying a specific aspect of the traces, and together allow for a detailed quantitative analysis of the experiment being analyzed. Specifically, the slope, rise time, and area measurements obtained from running the CSV_Peak_Detection notebook allow for quantitative analysis of the increase in cytosolic Ca^2+^ concentrations in cells exposed to trypsin and CCh.

### 3.3 SOCE Experiments

A further common experimental task is to quantify the content of intracellular Ca^2+^ stores and the extent of SOCE. The principal molecular components of SOCE are the ER Ca^2+^ sensors Stromal Interaction Molecule 1/2 (STIM1/2) and plasma membrane channels of the ORAI family [16]. The content of the stores is typically investigated by exposure of cells to a SERCA pump inhibitor such as cyclopiazonic acid (CPA) or thapsigargin in a bathing media which is depleted of Ca^2+^. The magnitude of the rise as Ca^2+^ leaks from the ER into the cytoplasm, together with the rate of rise, and area under the curve (AUC) provide information related to the level of the ER store content. In this paradigm, subsequent readmission of extracellular Ca^2+^ reveals the extent and rate of SOCE following ER store depletion, aggregation of STIM molecules and ORAI channel activation.

#### 3.3.1 Example: SOCE Experiment

When analyzing this type of experiment, it is important to measure the slope of the increase in [Ca^2+^] in the cytosol, which is related to the rate of SOCE. Therefore, it is critical to accurately identify the maximum point of the concentration curves. Thus, unlike in CSV_Peak_Detection, CSV_Peak_Detection_SOCE does not rely on peak detection to find the peak of the concentration curve, as the rise can sometimes be too gradual for a peak detection method to work reliably. Instead, the maximum point within the stimulation zone is used. The latency point is still identified by the point at which the traces go above a user adjustable threshold level.

As seen in **Figure 4**, the exposure of cells to CPA in calcium-free imaging buffer from 200-800 s causes an increase in cytosolic Ca^2+^ concentrations up to the point in which extracellular Ca^2+^ is depleted, at which point cytosolic Ca^2+^ concentrations gradually return to baseline levels. The subsequent exposure to CPA in Ca^2+^ imaging buffer, as seen from 900-1074 s in **Figure 4**, shows how a replenishment of extracellular Ca^2+^ again causes the cytosolic Ca^2+^ concentrations to rise. With no limitation of extracellular Ca^2+^ concentration, this concentration will not return to baseline before the end of the experiment. The slope and latency of both increases are important measurements in these experiments.

**Figure 4.**
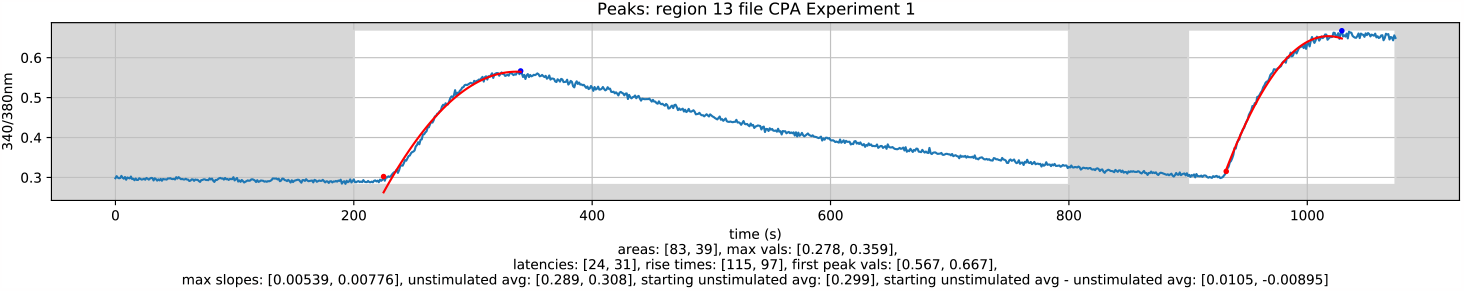
Example analysed plot output from CPA Experiment 1 Region 13

Note that, specifically, the rise time, slope, and maximum results from the CSV_Peak_Detection_SOCE notebook allow quantitative analysis of the extent to which the ATPases of the cell are inhibited. Using multiple stimulation zones (indicated with white background color in **Figure 4**, and automatic baselines also allows for the analysis of the extent to which the functionality of ATPases returns once no longer exposed to CPA.

### 3.4 Inhibitor Experiments

A frequent experimental question is the effect of an inhibitor of the Ca^2+^ signaling machinery. Here, as an example, we analyzed the effect of the ORAI inhibitor, GSK7975a [17], on the SOCE initiated by readmission of extracellular Ca^2+^ following exposure to CPA, as described in the previous section. The notebook is however designed to yield important parameters from a wide variety of experimental situations investigating the potency, efficacy, and kinetics of inhibitors of Ca^2+^ signaling.

### 3.4.1 Example: GSK Experiment

The latency and slope of the decrease in cytosolic Ca^2+^ concentration is an important measurement in experiments to assess the effects of inhibitors. In addition, the difference between the initial cytosolic Ca^2+^ concentrations and lower cytosolic Ca^2+^ concentrations following exposure is an important parameter.

Analysis begins with ROI identification, the details of which can be found in the supplemental notes. Analysis continues using the CSV_Peak_Detection_Inhibitor notebook. This notebook allows for analysis of antagonist experiments, which calls for negative latency thresholds and negative slope measurements. When analyzing inhibitor experiments, it is important to measure the slope of the decrease in concentration of Ca^2+^ in the cytosol.

Therefore, the minimum point within the stimulation zone is identified and used in the slope calculation. The latency point is determined as the point at which the trace goes below the calculated threshold level as opposed to above it with the other notebooks. The result is the ability to measure the downward slope of the trace.

Figure 5. shows a plot of one example region which includes the measurement of the slope of decrease in cytosolic Ca^2+^ concentration. Another stimulation zone could have been selected, say from 300-400 s, and the automatic baseline from this stimulation zone could have been compared to the starting baseline to measure the difference in initial cytosolic Ca^2+^ concentrations and lower cytosolic Ca^2+^ concentrations caused by GSK exposure.

**Figure 5.**
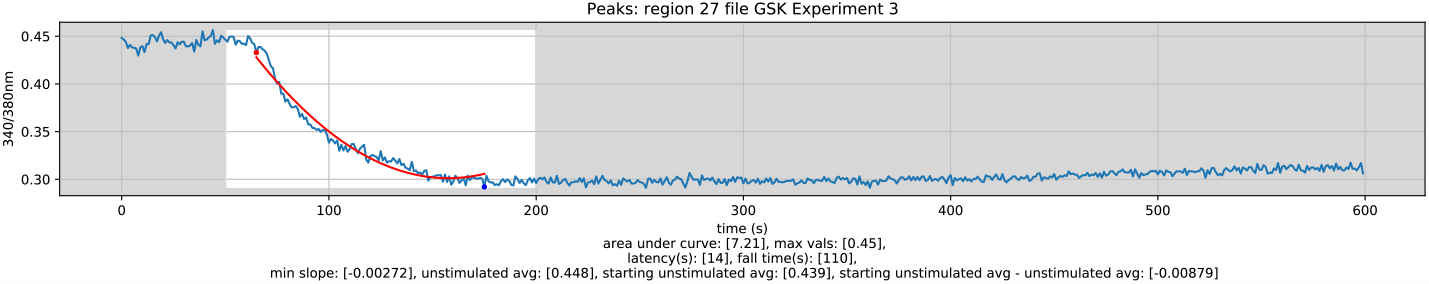
Example analysed plot output: GSK Experiment 3 Region 27

Note that, specifically, the slope and fall time measurements output by the CSV_Peak_Detection_Inhibitor notebook allow for a quantitative analysis of the effects of ORAI inhibition on cytosolic Ca^2+^ concentrations. By using multiple stimulation zones and automatic baselines, it is also possible to quantify the change in baseline Ca^2+^ levels in the cytosol as a result of ORAI inhibition.

## 4. Discussion and Conclusion

We found that the notebook paradigm is a good conceptual fit in a laboratory environment which contributes to ease of use.

Why didn’t we use the well-known software tool ImageJ [18]? In fact, we do use ImageJ for, among other things, the microscopy file format conversions and for initial exploration of raw image stack data. Yes, ImageJ targets microscopy image analysis, has a graphical user interface, supports scripting and packages together a very rich collection of analysis tools including generic region of interest selection. However, because we sought a lean user interface that only shows what is needed, has a self-documenting clear presentation of workflow and an ease of use that targets our specific application, we chose Python and Jupyter. With that said, the notebooks do not attempt to redo anything that is already done well with some other software tool. For example, the microscope manufacturer’s software tools are indispensable for data acquisition and for initial pre-processing operations, such as image deconvolution.

Other researchers, such as, for example, Leigh, et al. [19], have developed custom methods for identifying regions of interest in microscopy image stacks of Ca^2+^ signaling. However, in our case, because of a priori knowledge of cell size and shape, we were able to employ a very much less complex computational approach.

Initial interpretation of Ca^2+^ signaling data is often qualitative and at least somewhat subjective. However, given mathematically well-defined criteria and consistent parameters, repeatable analysis is possible. The resultant consistent quantitative analysis largely eliminates the subjective component.

In practice, we found that peak identification, as opposed to frequency analysis, is preferred by users for characterizing response plots. This is probably due the fact that peak locations are more immediately intuitively meaningful than are relatively abstract frequency plots.

We found that in general, our notebook collection, as a toolset and template, easily accommodates differing experiment requirements.

Because our Jupyter notebooks contain script source code in readable plain text, nothing is hidden, the analysis process is transparent. The notebook collection is available for use by other research groups as publicly hosted downloadable open-source code in a repository on GitHub [20]. Instructions for downloading the example datasets are included in the repository. The notebooks, along with the example data, can be used to reproduce the plots in this paper.

It does take some time to acquire a working familiarity with Python and Jupyter notebooks. However, both were designed with ease-of-use in mind and there are readily available online learning resources. We hope that other experimentalists will find these notebooks useful either as they are, or as a starting template for further bespoke customization. The learning curve for navigating and modifying Jupyter notebooks is modest but not trivial. In our experience the benefit far outweighs the “cost”.

## 5. Acknowledgements

New Zealand eScience Infrastructure (NeSI), for providing high-performance computing resources and consulting services.

University of Auckland, Centre for eResearch, for access to cloud computing resources and consulting services.

This work was supported by NIDCR grants 2R01DE019245-11 and F31 DE030670.

## Notes

https://github.com/jrugis/cell_tools/blob/master/Supplemental_Notes.pdf

### Competing Interest Statement

The authors have declared no competing interest.

